# Bioremediation of chlorpyrifos and 3,5,6-trichloro-2-pyridinol (TCP) by a paddy field bacterial isolate: Insights from genome analysis for possible biodegradation pathway

**DOI:** 10.1101/2019.12.12.866210

**Authors:** Tanmaya Nayak, Tapan Kumar Adhya, Ananta N Panda, Bhaskar Das, Vishakha Raina

## Abstract

Chlorpyrifos (CP) is one of the world’s most widely used organophosphorous pesticides (OPs) in agriculture, has led to contaminate soil and water ecosystems along with human species. Here, we report a newly isolated strain, *Ochrobactrum* sp. CPD-03 form a paddy field which could degrade CP and its metabolite TCP (3,5,6-trichloro-2-pyridinol) in 48 hours with an efficiency of 85-88% in minimal salt medium. GC-MS analysis revealed possible metabolites during CP degradation. Whole genome analysis indicated the presence of arylesterase and existence of other genes accountable for xenobiotic compounds degradation. CPD-03 exhibited chemotactic features towards CP along with other OPs suggesting its versatile role for possible mineralization of these toxicants. Further screening of CPD-03 also displayed plant growth promoting activities. Taken altogether, our results highlight the potentials of this new isolate *Ochrobactrum* sp. CPD-03 in bioremediation and application in OP-contaminated ecosystem.

## 1. Introduction

Current agriculture has been studied continuously using organic pesticides to produce improved crop yields and food health (Aswathi et al., 2019). While pesticides play an important role in the growth of crops in modern agriculture, widespread use does significant harm to the environment (Barbieri et al., 2019). The use of organophosphorus pesticides (OPs) have triggered alarm over environmental pollution, food safety and contact toxicity. OPs are the most toxic substances known and are widely used for pest control (Abhilash and Singh, 2009). Furthermore, release of pesticide residues into the atmosphere leads microbes to either detoxify these toxic molecules or use them as new sources of carbon and energy. Mineralization of these OPs is significantly more challenging since microbes need specific enzymes to turn such molecules into certain intermediates in a key metabolic pathway (Copley, 2009). However, given adequate exposure time, microbial cells may develop pathways to mineralize these pesticides (Lal et al., 2006).

Chlorpyrifos (CP; *O,O*-diethyl *O*-3,5,6-trichloropyridin-2-yl phosphorothioate) is considerably toxic (Mhadhbi and Beiras, 2012) among OPs commonly used for protecting agricultural crops from insects, fungi, weeds and other pests which affect the economic growth of crops. It enters inside the insects through ingestion, by contact or absorbed through cuticles, gut (Harishankar et al., 2013) and affects the nervous system by inhibiting the acetylcholinesterase activity, ultimately causing death of the target insect (Abreu-Villaca and Levin, 2017). CP hydrolysis leads to the formation of TCP (3,5,6-trichloro-2-pyridinol), a key toxic metabolite having considerable water solubility and antibacterial potential (Das and Adhya, 2015). TCP leaks into the soil and surface water bodies which cause extensive soil and aquatic habitat pollution, posing significant ecotoxicological threats owing to its neutral pH as compared to its parent compound and has a half-life between 65-360 days in soil (Li et al., 2010) and posing equal intimidations as CP. The enhanced resistance to microbial degradation of TCP is due to the existence of three chloride atoms in its chemical structure (Singh et al., 2003; Jabeen et al., 2015) and upon release of these inhibits the growth of the CP degrading microbes in the neighboring environment.

Living species being exposed to these CP residues present in soil and water, affects ecosystem balance (Chen et al., 2012). CP toxicity has been shown to be liable for nervous system disorders, immune system defects, liver damage and endocrine disruption. Hence, remediation of the CP contaminated sites to mitigate the effects of these toxic residues are highly important. Certain causes of pollution of this pesticide include industrial effluents, product waste, leakage, accidental spills and remediation. Therefore, techniques to degrade CP and TCP from contaminated fields need to be established in order to reduce the adverse effects on human health and ecosystem.

Bioremediation is one of the most promising methods due to its low cost and eco-friendly towards decontaminating pesticides (Rayu et al., 2012). Microbes containing the appropriate metabolic pathways play a key role in CP and TCP degradation (Singh et al., 2011). In the past, several bacteria such as *Enterobacter* sp. (Singh et al., 2004), *Bacillus* sp. (El-Helow et al., 2013), *Sphingomonas* sp. (Li et al., 2007), *Paracoccus* sp. (Xu et al., 2008), *Cupriavidus* sp. (Lu et al., 2013), *Brucella* sp., *Klebsiella* sp., and *Serratia marcescens* (Lakshmi et al., 2008), *Stenotrophomonas* sp. (Dubey and Fulekar, 2012) and *Stenotrophomonas* sp. (Yang et al., 2006), Alcaligenes (Yang et al., 2005), *Gordonia* & *Sphingobacterium* (Abraham et al., 2013), and *Mesorhizobium* (Jabeen et al., 2015) have been reported to utilize CP as a sole source of carbon. *Pseudomonas* sp. was the first ever bacterium reported for capable of utilizing TCP as a sole source of carbon and energy (Feng et al., 1997a) and another species, *Pseudomonas nitroreducens* AR-3 was reported for fastest CP degradation (Aswathi et al., 2019). *Ralstonia* sp. was studied for its TCP degradation ability (Li et al., 2010). While a large phylogenetic diversity of microbes has been established which can degrade OPs and their metabolites, biotic and abiotic environmental factors have a strong influence on the potential for degradation (Mrozik et al., 2010). Recently, *Ochrobactrum* sp. JAS2 was reported showing the ability of CP & TCP degradation in spiked soil within 24 h & 94 h respectively along with plant growth promoting ability (Abraham and Silambarasan, 2016). There are some instances where the bacterial species could lose its degradation ability under high pressure of TCP, thus making difficult for CP degradation in contaminated areas (Yadav et al., 2016).

The microbial strains showing degradation of pesticides had specific enzymes that exhibited particular action against xenobiotics. There are numerous reports concerning the efficacy of microbial enzymes in hydrolyzing chlorpyrifos (Gao et al., 2012; Fan et al., 2018). Aryldialkylphosphatases (EC 3.1.8.1), a group of hydrolytic enzymes, also known as organophosphate hydrolases or phosphotriesterases, show considerable activity against organophosphate triesters widely used as pesticides and phosphorofluoridates widely used in nerve agents (Raushel et al., 2000).

Organophosphate hydrolase (OPH) remains an important enzyme in chlorpyrifos detoxification which also contains methyl parathione hydrolase (MPH). Such enzymes hydrolyze a wide range of organic substrate such as paraoxone, parathione, diazinon and cyanophosis (Dumas et al., 1989). The OP degrading enzymes serve the same role but usually display little consistency between the enzyme sequences or pathways (Jiang et al., 2019).

The main objective of this study aimed to isolate, identify and select the active microbial strain(s) from paddy field soil capable for CP and TCP mineralization along with annotating a possible metabolic pathway responsible for CP degradation. This analysis would deepen our understanding and would provide valuable source of information to expand its potential as a biodegrader of OPs pesticides.

## 2. Materials & methods

### 2.1 Soil Samples, cultivation media and chemicals

Soil samples were collected from a paddy field near Kalinga Institute of Social Science [20.3551° N, 85.8187° E], Bhubaneswar Odisha, India. Preliminary screening of bacteria was performed using minimal salts media (MSM) [Per litre contains; NH_4_Cl, 0.5g/L; MgSO_4_.7H_2_O, 0.3g/L; KH_2_PO_4_.H_2_O, 0.2g/L; FeSO_4_.7H_2_O, 0.01g/L; Ca(NO_3_)_2_.4H_2_O, 0.01g/L;] supplemented with CP (100 mg L^-1^) as a soul carbon source. Analytical grade CP (PESTANAL^®^ 99.99%,) and TCP (PESTANAL^®^ 99.99%) were procured from Sigma-Aldrich, USA. Another commercial grade CP (TRICEL 20% EC and HILBAN 50% EC) were also purchased from local market for soil enrichment purpose. Chemicals and other reagents used in this study were of analytical grade (HiMedia, India and Merck, Germany).

### 2.2 Screening & isolation of CP degrading bacteria

CP degrading bacteria was isolated from rhizospheric soil by selective culture enrichment technique. Briefly 100 g of soil treated with CP (100 mg l^-1^) was inoculated into a 200 ml of MSM media. These soil samples were incubated at 30±2°C in a shaker incubator at 120 rpm for a week. Post incubation, 10 ml of grown culture was transferred for another week of incubation to new MSM amended with CP (200 mg L^-1^). Likewise, three consecutive transfers in sterile MSM supplemented with CP (100 mg L^-1^) was performed upon serially dilution and plated on solid media (MSM with CP @100 mg L^-1^). Post 96 h incubation, the pure colonies were obtained by repeated streaking. Isolate CPD-03 was further selected based on its CP degradation potential. Degradation efficiency was calculated [(Residual amount in blank control − residual amount in sample)/ (Residual amount in blank control) x 100]

### 2.3 Morphological, biochemical and taxonomic identification of the bacterial strain CPD-03

Isolate CPD-03 was preliminarily characterized by morphological parameters by scanning electron microscope (Nova™ NanoSEM 450) and Gram staining (HiMedia, India) followed by biochemical assays using BIOLOG^®^ microbial identification system (Engelen et al., 1998). Genomic DNA was extracted followed by a 16S rRNA sequencing (Weisburg et al., 1991) was performed. The 16S rRNA gene was obtained by PCR amplification with universal bacterial primers 27F and 1492R. PCR reaction was set at 95°C denaturation for 5 min, followed by 30 cycles of 95°C for 45s, 55°C for 45s, and 72°C for 90s, and a final step of 72°C for 5 min. The PCR amplified product was purified and sequenced by SciGenome, Ahmedabad, India. The obtained 16s rRNA gene sequence was aligned with the 16S rRNA sequences of type strains with homology of 97% or higher with CPD-03 and were obtained from EzTaxon (eztaxon-e.ezbiocloud.net). The sequence alignment was performed using ClustalX (Larkin et al., 2007). Phylogenetic tree was constructed with MEGA version 7.0 (Kumar et al., 2016).

### 2.4 Inoculum preparation (Resting cell assay)

Pure cultures of bacterial strain CPD-03 were cryopreserved in 25% glycerol at - 80^0^C. The bacterial strain was revived and inoculated in a 100 ml Erlenmeyer flask with 20 ml sterile MSM supplemented with CP (100 mg L^-1^) before each set of experiments. Flask was incubated at 30 ± 2°C with 120 rpm in a shaker incubator. The bacterial cells were harvested after 44 - 48h at 4500 rpm for 5 min followed by washing with sterile saline (0.9% NaCl). The washed bacterial cells were resuspended in sterile MSM to obtain a cell density of 1-1.5×10^8^ cells ml^-1^. This bacterial resting cell suspension was used for further biodegradation experiments, organophosphorus hydrolase (*opd*) gene localization, chemotaxis assays and plant growth promoting response (PGPR) activities.

### 2.5 Biodegradation of CP & TCP in aqueous medium

Biodegradation of CP (100 mg L^-1^) & TCP (100 mg L^-1^) was studied (in separate and in mixed) in 50 ml resting cell suspension (Supplementary Figure 1a&b). An additional carbon source molasses (0.1% v/v) was added to enhance the growth. MSM with the same concentration of CP and 0.1% molasses was treated as control. All the experimental flasks were kept for incubation at 30±2°C and 120 rpm. Degradation under various concentration of CP (100 mg L^-1^ to 1000 mg L^-1^) was studied in 50 ml resting cell as mentioned above were monitored till 12 h in an interval of 2 h and later till 48h in an interval of 6 h for determining the rate of CP degradation using high performance liquid chromatography (HPLC) (Agilent Technology 1260 Infinity, USA).

**Figure 1.**
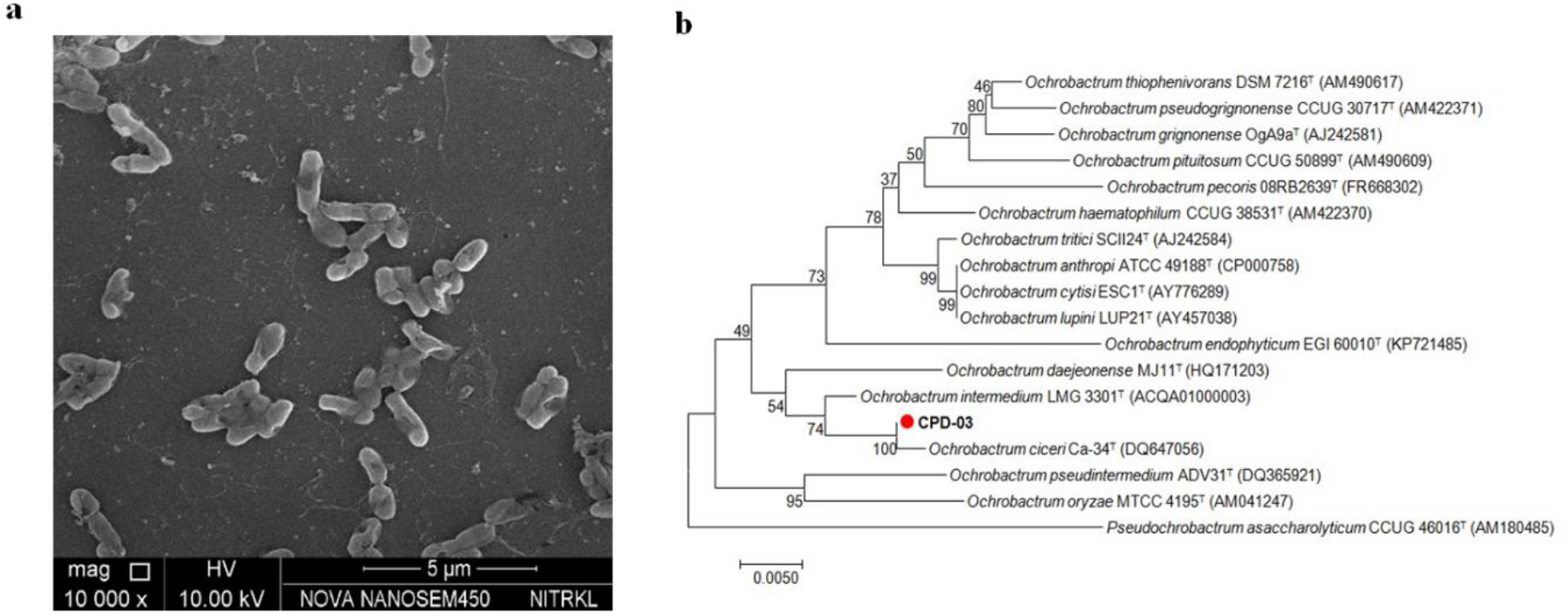
a. Scanning electron microscopy of CPD-03; b. phylogenetic tree of CPD-03 based on comparison of 16S rRNA sequence similarity of closely related *Ochrobactrum* sp. conducted in MEGA version 7.0

### 2.6 Sample extraction and HPLC analysis

Upon completion of biodegradation assay, the entire content including cells and residual CP, was harvested at 7500 rpm for 10 min for obtaining a cell-free media and extracted with organic solvents. The supernatant was pulled out for extraction with an equal volume of ethyl acetate and this step was repeated twice. The organic layer thus obtained was aspirated, pooled and dehydrated over a column of anhydrous sodium sulfate, evaporated at 45±2°C using rotavapor (Eyela, Japan), reconstituted in methanol (HPLC Grade) and filtered through 0.22 μm membrane filter. Residual CP and TCP was quantified using a reverse-phase C18 column (ZORBAX Eclipse Plus, Agilent Technologies, USA) fitted with Agilent Technologies 1260 Infinity HPLC system equipped with a binary pump, UV/VIS detector, thermostat column compartment (TCC) column oven and auto sampler, with array detection based on peak area and retention time of the pure standard. The mobile phase used for the analysis contained a mixture of methanol and water in a ratio of 85:15 at a flow rate of 1.2 mL min^-1^. Water (HPLC grade, Merck) was acidified to pH 3.2 using orthophosphoric acid. Injection volume was 20 µL. The recoveries of OPs added at concentrations of 1, 2, 5, 10, 50 mg/L, ranged from 97.0% to 100.0%. The linear range of the calibration curves was obtained from 1 to 50 mg/L.

### 2.7 Degradation kinetics

CP degradation rates has been fitted with a pseudo first order kinetic model *Y*= (*Y*_*0*_ - Plateau)*e^(-K*X)^ + Plateau; *Y*_*0*_ = the *Y* value when X (time) is zero and is expressed in the same units as *Y*, Plateau = the *Y* value at infinite times and expressed in the same units as *Y*. K = rate constant and expressed in reciprocal of the X axis time units. Half-life is in the time units of the X axis. It is computed as ln(2)/K using statistical software Prism8.

### 2.8 Identification of metabolites

The cell free extract was assessed for the identification of CP degradation and simultaneous metabolites formation. Samples were analysed in gas chromatography-mass spectrometry (GC-MS) (Agilent Technologies, USA) fitted with a HP-5MS capillary column (30.0 m x 250 µm x 0.25 µm) in mass spectrometer (Agilent 7890B ION TRAP MS) equipped with a manual injector containing split/split less capillary injection system along with an array detection of total scan from 30-500 nm. The column temperature was kept for 5 min at 80^0^C initially and increased for 5 min at 8^0^C min-1 to 200^0^C, then increased for 5 min at 15^0^C min^-1^ to 260^0^C. The ionization energy was kept at 70 eV, and the temperatures of the transfer line and the ion trap were kept 280^0^C and 230^0^C, respectively. The amount of sample injection was 1.0 μL at 250^0^C with 1:50 split mode. Helium was used as a carrier gas at a flow rate of 1.5 ml min^-1^. The CP and intermediates found by the GC-MS study matched authentic standard compounds from the National Standards and Technology Institute (NIST, USA) database (http://chemdata.nist.gov).

### 2.9 Molecular identification & localization of organophosphorus hydrolase (opd) gene

For identification of organophosphorus hydrolase (*opd*) gene, the resting cell (1-1.5×10^8^ cells ml^-1^) was used for genomic DNA isolation as mentioned section 2.3. The *opd* gene of strain CPD-03 was amplified by PCR with forward primer (OPDF, 5′-GC<u>GGATCC</u>ATGCAAACGAGAAGGGTTGTGC-3′, and reverse primer OPDR, 5′-GC<u>CTCGAG</u>TCATGACGCCCGCAAGGTC-3′; *BamHI* and *XhoI* restriction sites, respectively, are underlined). The reagents used for PCR were obtained from Promega (Madison, WI) and a total volume of 25 μl reaction mixture for PCR amplification was carried out. Each reaction mixture consists of 2× green reaction mixture (10 mM MgCl_2_, 10 μM each dNTPs, 1 U of Taq DNA polymerase along with 10x buffer), 10 pmol of each primer along with the template DNA and nuclease free water for volume makeup. The PCR amplification program was as follows: initial denaturation at 94°C for 5 min, 32 cycles of cyclic denaturation at 94°C for 60s followed by annealing at 55.1°C for 45s and extension at 72°C for 60s including a final extension at 72°C for 5 min.

To localize the presence of opd protein, cell-free supernatant was processed for enzyme assays (Yashphe et al., 1990). Briefly, CPD-03 cells were harvested at 8500×g for 15 in 4^0^C. Cell-free extract was obtained by passing the supernatant through a 0.22µm membrane filter and hence treated as extracellular (EC) and stored at 4^0^C. The cell pellet was washed twice with normal saline in the Tris-Cl buffer solution (50 mmol / L, pH 8.0) and re-suspended to obtain cell suspension (CS). One half of the CS was stored at 4^0^C, and the other half was re-suspended in Tris-Cl buffer and sonicated (4^0^C, 400W) for 8min. The collected suspension was then centrifuged at 8000 ×g for 10 min in 4^0^C. The resulting supernatant was filtered through a 0.22μm to obtain crude intracellular (IC) enzyme. *Opd* activity was calculated as the number of moles of *p*-nitrophenol produced in unit time.

### 2.10 Assessment of plant growth promoting activity

Evaluation of the effect of CPD-03 resting cell (1-1.5×10^8^ cells ml^-1^) on plant growth was done by quantitative measurement of root length in inoculated rice seedlings followed by indole acetic acid (IAA) production (Bric et al., 1991), ammonia production and HCN production (Bakker and Schippers, 1987). The data were subjected to statistical analysis using Prism8.

### 2.11 Chemotaxis towards CP

The chemotactic property of CPD-03 resting cell (1-1.5×10^8^ cells ml^-1^) towards CP along with other OPs was examined qualitatively in swarm plate assays and quantitatively with capillary tube assay as per the procedure described earlier (Pandey et al., 2012). Citrate (0.5%) was used as the positive control (Pailan and Saha, 2015). The chemotactic response was expressed as chemotaxis index (CI), and calculated as the ratio of the number of CFUs obtained from the capillary tube containing the test compound (CP) to CFUs obtained from a control capillary that contains only the chemotaxis buffer (100 mM potassium phosphate pH 7.0 and 20 μM EDTA; without any chemotactic compound).

### 2.12 De novo genome sequencing and assembly

Illumina HiSeq System 2000 (Illumina, Inc.) was used for whole-genome sequencing. Data obtained from sequencing were *de novo* assembled followed by entire quality control performed using NGSQC Toolkit. Velvet (V. 1.2.10) (Zerbino and Birney, 2008) was used for primary assembly that resulted a total size of the contig of 4,65,9698 (∼4.6Mb). *De novo* Genome Validation was performed by Bowtie2 (v 2.2.2) (Langmead and Salzberg, 2012). Quality control of assembled genome based on genomic elements were performed by ARAGORN (v1.2.36) (Laslett and Canback, 2004). Genome Annotation and functional characterization were performed using RAST (http://rast.nmpdr.org/). All the bio-informatics work was performed in collaboration with Bionivid Technology Pvt. Ltd., Bangalore, India. Genome sequence including the necessary information for the CPD-03 strain was deposited in NCBI under the accession number RSEU00000000. Considering the ability of this strain for degrading CP, the whole genome of this strain was checked for the occurrence of the CP degrading core genes/proteins.

## 3 Results and discussion

### 3.1 Isolation and identification of CP degrading bacterial strain CPD-03

Several bacterial strains were isolated from rhizosphere soil of paddy fields near KIIT University area which could utilize CP as carbon source for their growth. CPD-03 was distinguished from other isolates for its ideal growth response during isolation. Further CPD-03 (1.2×10^8^ cfu/ml) showed a CP degradation efficiency of 82±2% in MSM supplemented with CP (100 mg L^-1^) (Supplementary Figure.1a) and hence selected for further study. CPD-03 was able to degrade TCP (100 mg L^-1^) separately with a degradation efficiency of 60±2% (Supplementary Figure.1b).

CPD-03 is Gram-negative and showed regular rod-shaped cells which was observed in scanning electron microscopy (Figure. 1a). A detailed comparison of the physiological and biochemical properties of CPD-03 with its closest phylogenetic species of *Ochrobactrum* was performed (Supplementary Table 1a-c). A phylogenetic tree including strain CPD-03 and related type strains within the clade of *Ochrobactrum* (obtained from EzBioCloud) was also compared (Figure. 1b). 16S rRNA gene sequence analysis illustrated that, CPD-03 shared maximum homology of 99.8% with *Ochrobactrum intermedium* LMG 3301^T^ followed by 99.2% with *Ochrobactrum ciceri* Ca-34^T^ of the genus *Ochrobactrum* (Imran et al., 2010), hence designated as *Ochrobactrum* sp. CPD-03. The 1396 bp 16S rRNA nucleotide sequence of CPD-03 strain was submitted in GenBank under Accession No. KT366923.

### 3.2 Biodegradation of CP and TCP

Preliminary CP degradation studies by CPD-03 was earlier monitored in lab (Nayak et al., 2020). Followed to this, degradation of CP (100 mg L^-1^) and TCP (100 mg L^-1^) were assessed using CPD-03 resting cell (1-1.5×10^8^ cells ml^-1^) in MSM along with molasses (0.1% v/v). This resting cell approach showed CP and TCP degradation efficiency of 88±2% and 62±2%, respectively in 48 h (Figure. 2a). TCP degradation was monitored separately using CPD-03 resting cell (1-1.5×10^8^ cells ml^-1^) in MSM along with molasses (0.1% v/v) with a degradation efficiency of 62.2±2% (Figure. 2b). Earlier reports have shown TCP can be degraded by *Alcaligenes* sp. (Yang et al., 2005), *Burkholderia* sp. (Kim and Ahn, 2009), *Pseudomonas* sp. (Feng et al., 1997b), *Pseudomonas* sp., *Bacillus* sp. and *Agrobacterium* sp. (Maya et al., 2011), *Trichosporon* sp. (Xu et al., 2007), and *Pseudomonas* sp. (Feng et al., 1997a) when used as sole carbon source. Previous report has shown *Ochrobactrum* sp. JAS2 had degraded not only CP and TCP in aqueous medium within 96 h when used together. The present study showed, CPD-03 could degrade CP at various concentrations (100 mg L^-1^ to 1000 mg L^-1^) and it was clearly evident that higher concentration reduced the CP degradation efficiency (Figure. 2c). This correlates with a previous report where higher concentration of pesticides has inhibited the bacterial growth hence decreasing the degradation rate and highlights as the second report from the genus *Ochrobactrum* possessing the ability of CP and TCP degradation.

**Figure 2.**
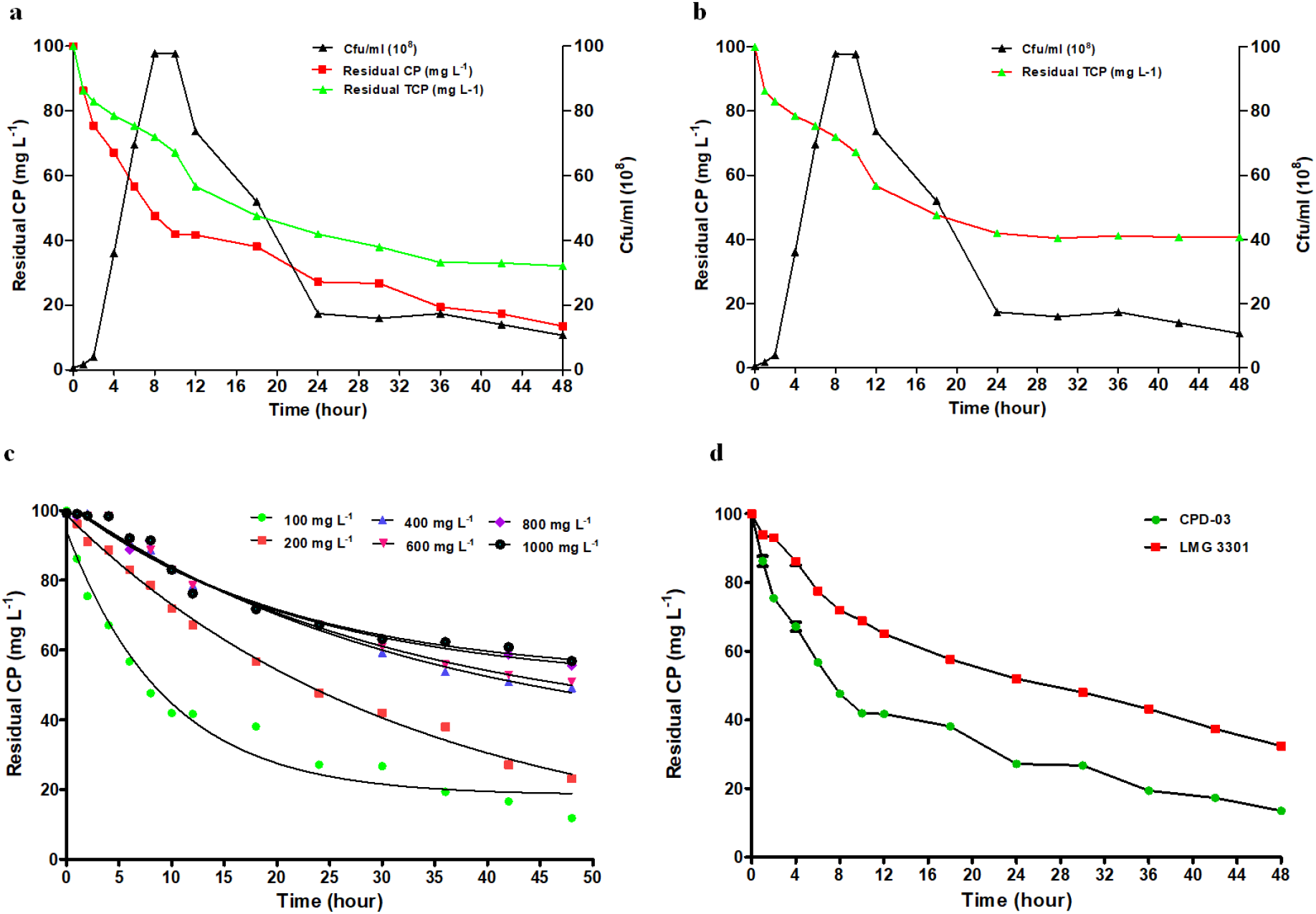
a. Degradation of CP (100 mg L^-1^) and TCP (100 mg L^-1^) by CPD-03 resting cell (1-1.5×10^8^ cells ml^-1^) in a MSM supplemented molasses (0.1% v/v) showing formation of its intermediate TCP and its further degradation; b. degradation of TCP (100 mg L^-1^) by CPD-03 resting cell (1-1.5×10^8^ cells ml^-1^) in a MSM supplemented molasses (0.1% v/v); c. degradation of CP at various concentrations (100 mg L^-1^, 200 mg L^-1^,400 mg L^-1^,600 mg L^-1^,800 mg L^-1^, and 1000 mg L^-1^) by CPD-03 resting cell (1-1.5×10^8^ cells ml^-1^) in MSM supplemented molasses (0.1% v/v); d. comparision of CP (100 mg L^-1^) degradation by CPD-03 resting cells (1-1.5×10^8^ cells ml^-1^) and LMG 3301^T^ (1.2-1.4×10^8^ cells ml^-1^). Data were represented in the mean of three replicates and analysis were performed using ANOVA with the Prism8. All were tested at the *p<*0.005*** significance level.

CPD-03 resting cell in MSM supplemented with molasses (0.1%) revealed significant compliance for CP degradation (100 mg L^-1^ to 1000 mg L^-1^), was curve fit into a pseudo-first order kinetics [*Y*= (*Y*_*0*_ - Plateau)*e^(-K*X)^ + Plateau)] (Affam and Chaudhuri, 2013). A recent study had reported the CP degradation by *Ochrobactrum* sp. JAS2 also followed a pseudo first order kinetics (Abraham and Silambarasan, 2016). CPD-03 degraded 100 mg L^-1^ CP with a half-life of 6.559 h and TCP 100 mg L^-1^ with a half-life of 12.08 h. CPD-03 was compared with its phylogenetically closest neighbor *Ochrobactrum intermedium* LMG 3301^T^ for its ability to degrade CP (100 mg L^-1^) and was found to show higher degradation efficiency of 88.34% as compared to LMG 3301^T^ with a degradation efficiency of 60.34% (Figure. 2d).

### 3.3 Identification of possible metabolites

Degradation of CP by *Ochrobactrum* sp. CPD-03 was monitored in GC-MS, has resulted new metabolites (Figure. 3). Previous studies have shown the formation of TCP upon CP mineralization and its immediate degradation upon subsequent hydrolysis steps through ring cleavage (Chen et al., 2012). However, in our study, few new downstream metabolites like (a) cis-Vaccenic acid, (b) Octadecanoic acid, (c) Benzoic acid, 4-hydroxyl 3-methoxy, (d) n-Hexadecanoic acid, (e) 9-Hexadecanoic acid were obtained after CP mineralization which could add possible insights into the metabolic pathways. cis-Vaccenic acid is a byproduct from the activity of 3-oxoacyl-[acyl-carrier protein] reductase which was present in CPD-03 genome. The presence of this enzyme has previously been shown in case of poly aromatic hydrocarbon (PAH) degradation (Lyu et al., 2014). Similarly, Octadecanoic acid was found to be a metabolite during biodegradation of endocrine-disrupting chemicals in earlier reports (Chandra et al., 2018), Benzoic acid, 4-hydroxyl 3-methoxy was reported in beta-cypermethrin degradation (Chen et al., 2013), n-Hexadecanoic acid production and degradation due to microbial activity has been studied (Zyakun et al., 2011) and 9-Hexadecanoic acid was reported to produce during bioremediation of *n*-alkanes (de Carvalho, 2012). This could possibly draw some new insights into the metabolic pathway in terms of pesticide degradation.

**Figure 3.**
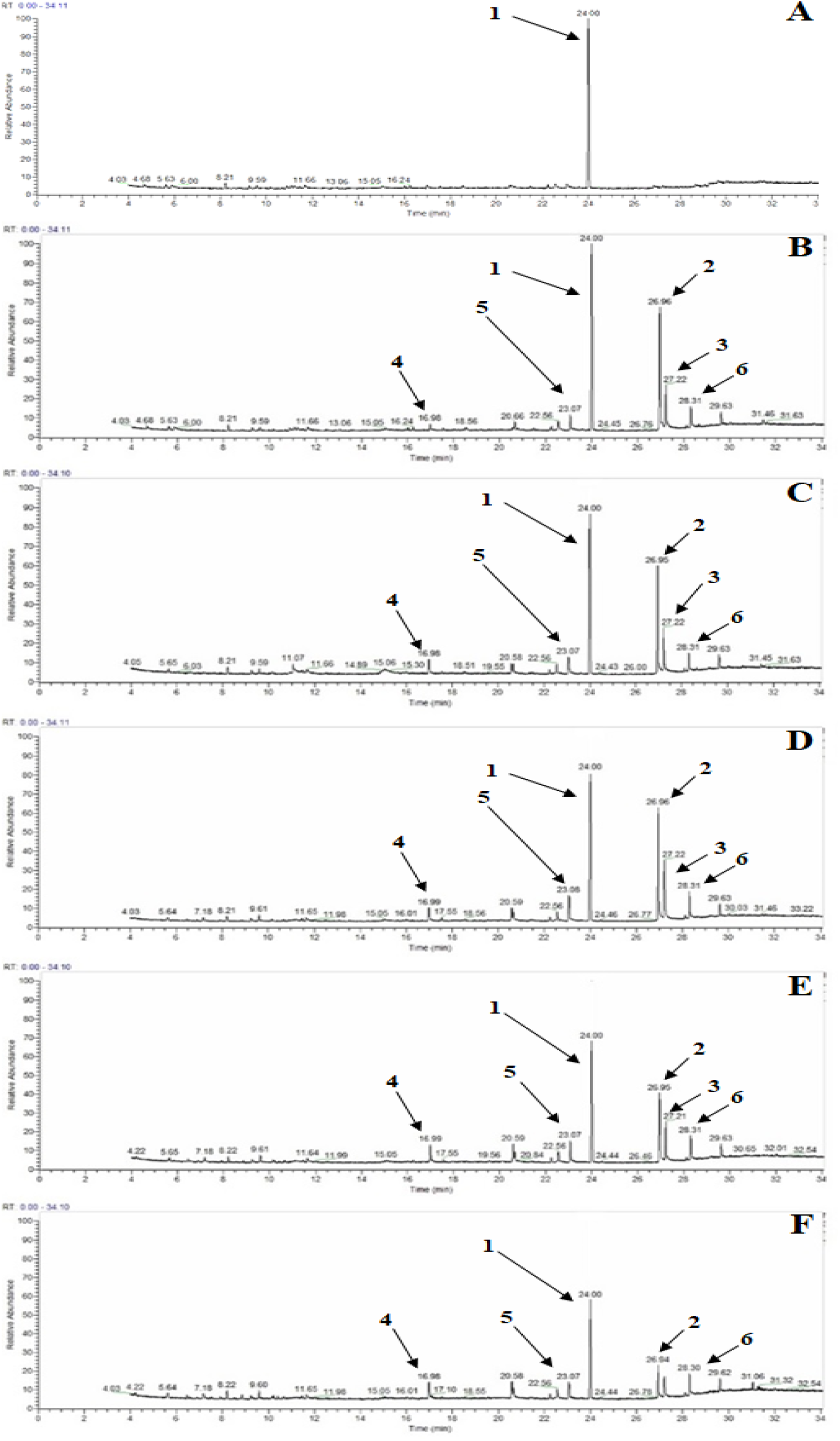
GC-MS chromatograms showing the metabolites formed during CP (100 mg L^-1^) degradation by CPD-03 resting cells (1-1.5×10^8^ cells ml^-1^). A: 0 h, B: 2 h, C: 4 h, D: 6 h, E: 8 h, and F: 10 h respectively. 1: chlorpyrifos, 2: cis-Vaccenic acid, 3: Octadecanoic acid, 4: Benzoic acid, 4-hydroxyl 3-methoxy, 5: n-Hexadecanoic acid, 6: 9-Hexadecanoic acid, respectively.

### 3.4 Localization of opd gene in CPD-03

CPD-03 was found to contain the gene encoding organophosphorus hydrolase (*opd*, ∼1098 bp), which was successfully amplified using the primers OPDF and OPDR (Figure. 4a). This genes could possibly transform the complex OPs to generate *p*-nitrophenol have been previously reported (Tang et al., 2014). Since CPD-03 is a Gram-negative bacteria, organophosphorus hydrolases potentially are secreted into the extracellular space from periplasmic region, hence can be extracted in more quantities (Wang et al., 2009). A maximum amount of *p*-nitrophenol (2±0.2 mmol/L) produced by treatment of the CPD-03 resting cells (extracellular cell free content, EC) with CP for 30 min (Figure. 4B) as compared to the intracellular and cell suspension. This result is consistent with the earlier report of *Flavobacterium* sp. MTCC 2495 (Falahati-Pour et al., 2016).

**Figure 4.**
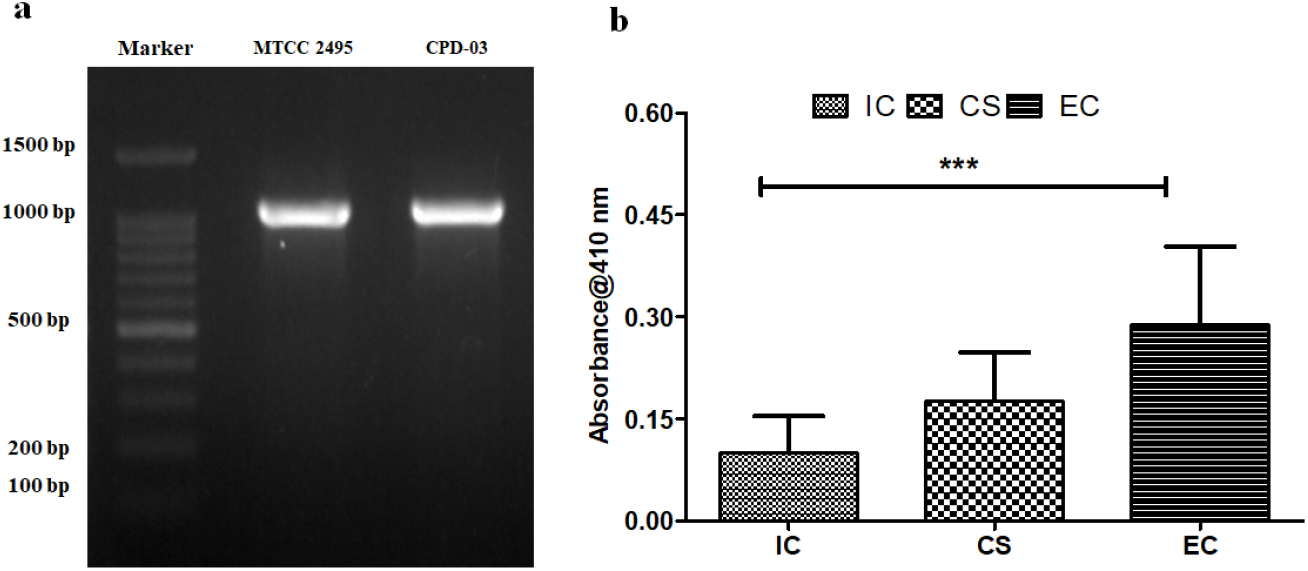
a. Amplification of ∼1098 bp *opd* gene; b. enzyme activity of CPD-03 resting cell (1-1.5×10^8^ cells ml^-1^); where IC: intracellular, CS: cell suspension and EC: Extracellular. Data analysis were performed using ANOVA with the Prism8. All were tested at the *p<*0.005*** significance level

### 3.5 CPD-03 genome analysis and presence of arylesterases gene

A comparative genomics approached was analyzed using KAAS (KEGG Automatic Annotation Server) (Moriya et al., 2007) for functional annotation of proteins and metabolic pathways present in CPD-03. The minimal sets of metabolic pathways were further annotated and reconstructed against the predicted protein families using a web-based tool MinPath (Ye and Doak, 2009) and were clustered using ClustVis (Metsalu and Vilo, 2015). In addition, comparative genome analysis of other *Ochrobactrum* genomes were assessed for the capability to degrade the aromatic and relative compounds (Supplementary Figure. 2). This feature could also demonstrate the potential genes of CPD-03 strain for possible application in case of aromatic compound degradation. Although microorganisms have been known to possess numerous enzymes with capability to degrade several xenobiotic compounds, only ten organophosphorus hydrolase genes were reported to degrade CP. Organophosphorus hydrolase (OPH; EC 3.1.8.1) is an ortholog of arylesterase (EC 3.1.1.2), are known to catalyze the hydrolysis of a wide range of OPs and are produced by few microbes. Due to its bioremediation potential, OPH has been used in industrial, health care and military purposes (Jackson et al., 2014).

A phylogenetic tree was constructed between the arylesterase sequence present in CPD-03 along with other phylogenetically close *Ochrobactrum* genomes. The highest sequence homology was obtained with *Ochrobactrum intermedium* LMG 3301^T^ (Supplementary Figure. 3). Additionally, comparison of *opd* gene sequence obtained from whole genome of CPD-03 and amplified *opd* gene sequence validated the sequence similarities in genetic and protein level (Supplementary Figure. 4). CPD-03 shared numerous putative pathways for aerobic degradation of aromatic compounds like benzoate, aminobenzoate, chloroalkane and chloroalkene, chlorocyclohexane and chlorobenzene (Supplementary Table 2) suggesting that it might have acquired genes and pathways for mineralization of these xenobiotic compounds from the soil ecosystem. Based on the previous reports along with the CPD-03 genome, a metabolic pathway has been proposed (Supplementary Figure. 5) which indicated the presence of gene clusters responsible for CP degradation by CPD-03.

**Figure 5.**
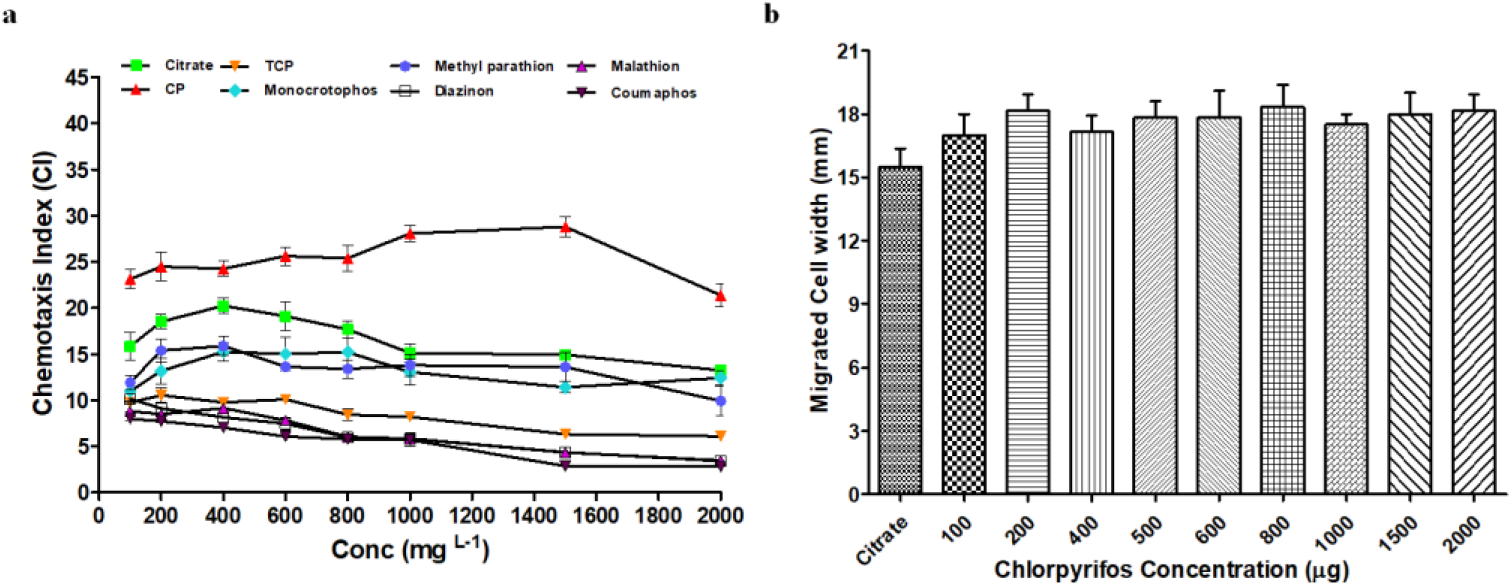
a. Chemotaxis index of CPD-03 resting cells CPD-03 (1-1.5×10^8^ cells ml^-1^) towards CP, TCP, monocrotophos, methyl parathion, coumaphos, diazione and malathione as competitive attractant at various concentrations (from 200 mg L^-1^ to 2000 mg L^-1^). Values are presented as arithmetic means and error bars indicate standard deviations based on three independent replicates. b. migrated cell width of chemotactic movement by CPD-03 towards various concentration of CP (100 mg L^-1^, 200 mg L^-1^, 400 mg L^-1^, 600 mg L^-1^, 800 mg L^-1^, 1000 mg L^-1^, 1500 mg L^-1^ and 2000 mg L^-1^) along with citrate (0.5%) as positive control. Data analysis were performed using ANOVA with the Prism8. All were tested at the *p<*0.005*** significance level.

### 3.6 Chemotactic response of strain CPD-03 towards various OPs

Bacterial chemotactic movement under a chemoattractant helps bacterium find metabolizable substrates for growth and survival. The chemoattractant is often a compound that also acts as a source of carbon and energy (Pandey and Jain, 2002) and has recently become a widespread phenomenon of bacterial metabolism against various xenobiotic compounds, and the use of these bacteria in bioremediation may therefore be beneficial (Gordillo et al., 2007). Chemotaxis ability of CPD-03 was studied against CP along with various other OPs such as monocrotophos, methyl parathion, diazinon, malathion, and coumaphos. The strongest chemotactic response was observed for CP as compared to other OPs (Figure. 5a). Moreover, various concentration of CP also had the similar chemotaxis phenomenon (Figure. 5b, Supplementary Figure. 6). Additionally, putative gene clusters were also found in the genome of *Ochrobactrum* CPD-03, validating the chemotactic features (Supplementary Table 3). To the best of our knowledge, our results suggested the first report of the genus *Ochrobactrum* for chemotaxis towards CP along with other OPs.

**Figure 6.**
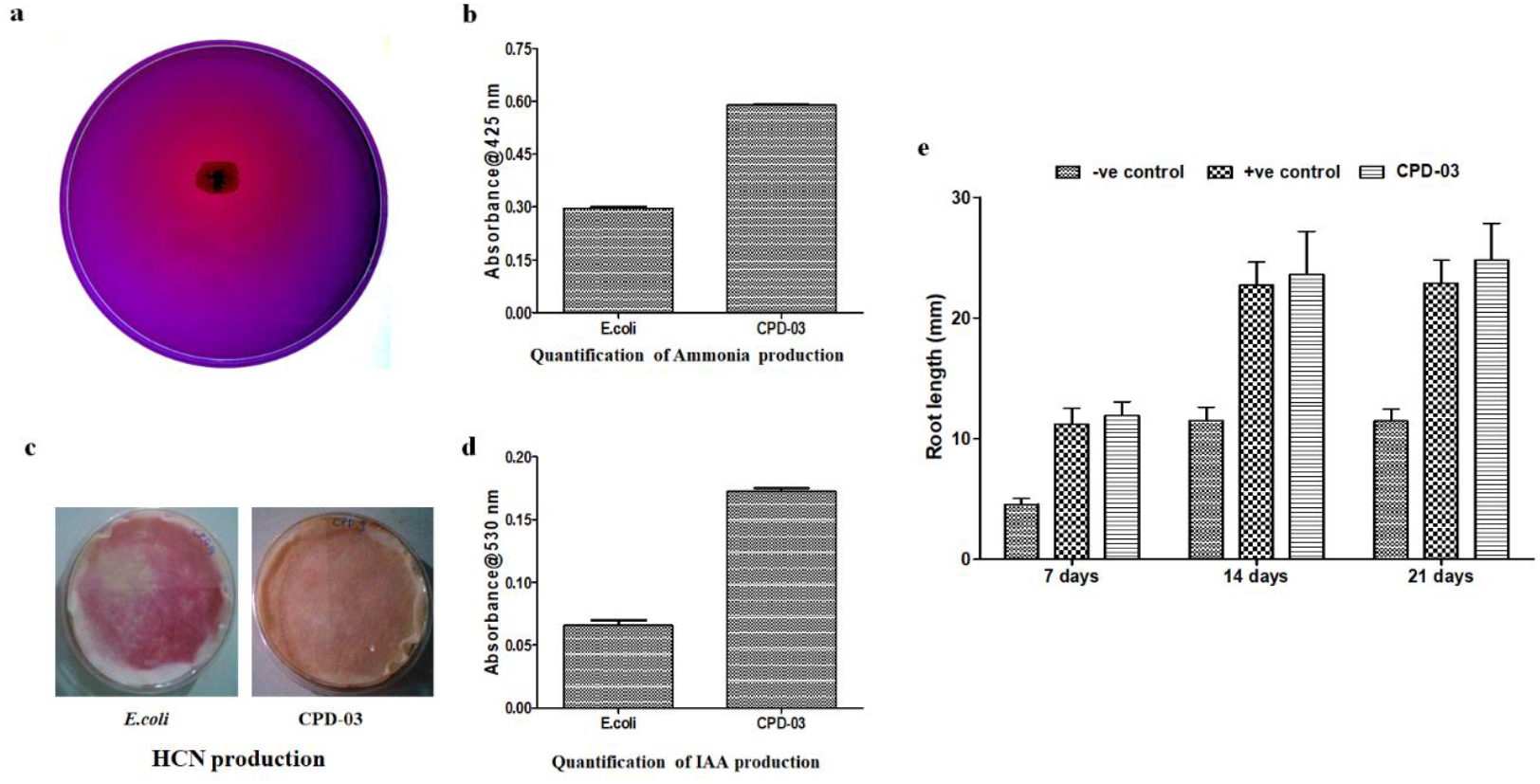
**(a-e)**. a. Phosphate solubilization on Pikovskaya’s agar supplemented with tricalciumphosphate by CPD-03 resting cells (1-1.5×10^8^ cells ml^-1^); b. quantification of ammonia production; c. HCN production and d. quantification of IAA production by CPD-03 resting cells (1-1.5×10^8^ cells ml^-1^); e. Effect of root length upon CPD-03 inoculation on rice seed; -ve control: culture free, +ve control: *Bacillus* sp., lab stock for positive PGPR activity. *E.coli* was used as a negative control. Data analysis were performed using ANOVA with the Prism8. All were tested at the *p<*0.005*** significance level

### 3.7 Assessment of Plant growth promoting ability of CPD-03

Rhizospheric bacteria live on and around the paddy fields have been contributing towards degradation of toxic xenobiotic compounds in contaminated ecosystem and shown to have potential for accelerating degradation process. Paddy field environment comprised of phosphorus in soils (in both organic and inorganic forms) (Khan et al., 2009). As a result, microbes associated with phosphate solubilizing ability would give an insight into the available forms of phosphate to the plants. In our present study, CPD-03 was found to solubilize the tricalcium phosphate post incubation, which was a qualitative analysis i.e. the agar media of the pikovskaya containing bromophenol turned from blue to yellow due to a decrease in the pH of the medium confirming a positive feature of phosphate solubilization (Figure. 6a). In plants, production of ammonia is accountable for the indirect plant growth promotion via pathogens control and d HCN is a key compound during this metabolism (Blumer and Haas, 2000). CPD-03 showed a substantial increase in ammonia production as compared to the control (*E.coli*) (Figure. 6b). CPD-03 had shown to produce HCN (Figure. 6c). It is reported that, microbes isolated from various agricultural sites have the ability to produce phytohormone auxin indole-3-acetic acid (IAA) as secondary metabolites (Patten and Glick, 1996). Generally, IAA secreted by microbes stimulate seed germination followed by increase the rate of xylem and root development, controls vegetative growth process, regulates photosynthesis, formation of pigments, production of several metabolites, and resistance to stress (Ahemad and Kibret, 2014). In our study, CPD-03 produced 40.33 mg L^-1^ IAA as compared to the control (*E.coli*) which had produced 22.36 mg L^-1^ of IAA, respectively (Figure. 6d). CPD-03 inoculation was also positively affected the increased root length (Figure. 6e). This result is consistent with the previous report on *Ochrobactrum* sp. JAS2. Genome analysis had shown putative gene clusters present in *Ochrobactrum* CPD-03, validating the potential effects of PGPR activity. However, the gene clusters for HCN production, phosphate solubilization, ACC deaminase activity, and ammonia production were not found. The genome shows the presence of several hypothetical proteins which might be contributing for these above mechanisms or presence of ortholog sequences (Supplementary Table 3) which has still not been studied but may be of interest in future.

## 4. Conclusions

CP degradation by *Ochrobactrum* sp. CPD-03 was isolated from paddy field with additional chemotactic ability towards other OPs and plant growth promoting abilities. Whole genome analysis of CPD-03 revealed a possible metabolic pathway the presence of gene clusters responsible for CP biodegradation. This study provides the genetic information for its application for the degradation of OP pollutants in natural environment for further development of a biostimulation process. Moreover, the plant growth promoting abilities could be of great potential for improved and persistent crop production during CP induced stressed agricultural ecosystem.

## 5. Acknowledgement

This research work was funded by the Department of Biotechnology (DBT), Govt. of India, New Delhi. Research grant sanction no. BT/PR7580/BCE/8/1009/2013. All authors remain highly grateful to the Director of the institute for infrastructure that enabled us to perform this work.

