## Supplementary Information for "Bioremediation of chlorpyrifos and 3,5,6-trichloro-2-pyridinol (TCP) by a paddy field bacterial isolate: Insights from genome analysis for possible biodegradation pathway"

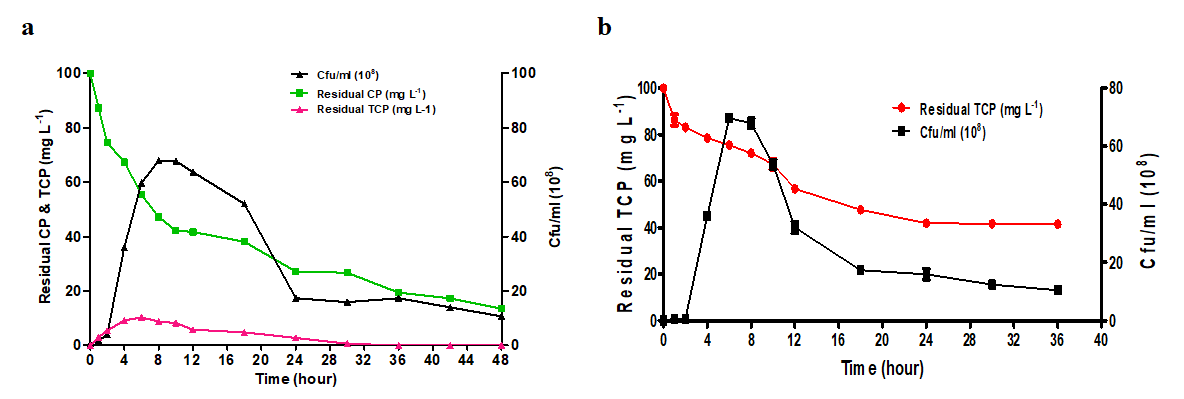

**Supplementary Figure.1** a. Degradation of CP (100 mg L^-1^) by CPD-03 (1.2x10^8^ cfu/ml) leads to formation of TCP followed by its further degradation; b. degradation of TCP (100 mg L^-1^) by CPD-03 (1.2x10^8^ cfu/ml). Data were represented in the mean of three replicates and analysis were performed using ANOVA with the Prism8. All were tested at the *p<*0.0001*** significance level

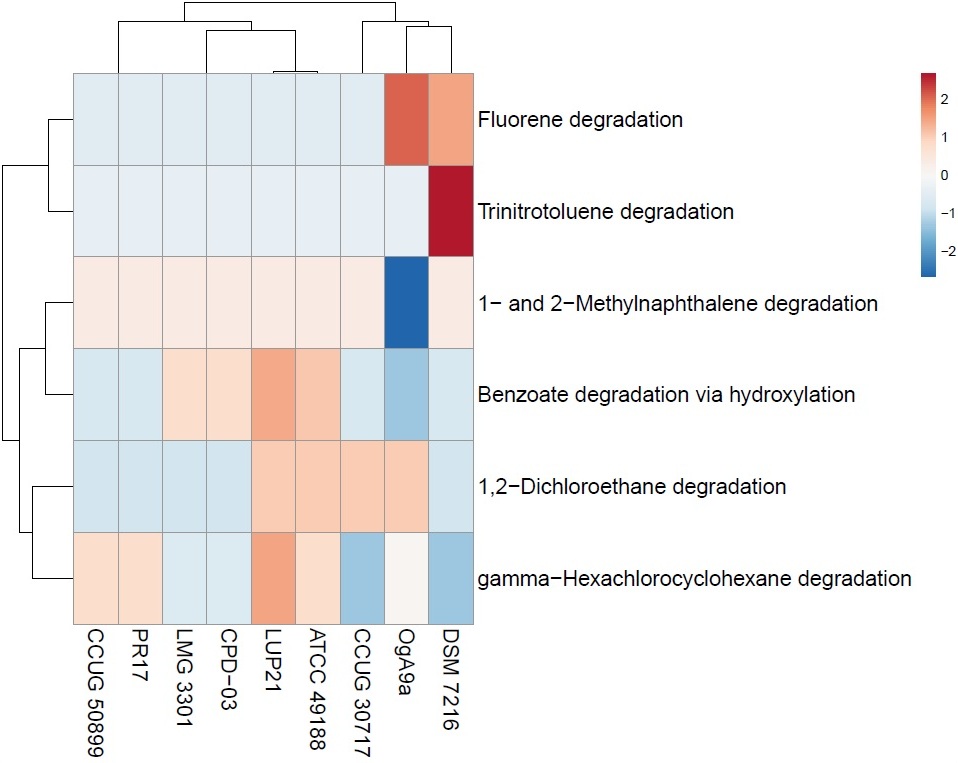

**Supplementary Figure. 2**. Heatmap and dendrogram of potential abilities of degradation of Aromatic and relative compounds predicted by genomic features of CPD-03 along with other *Ochrobactrum* strains (*Ochrobactrum* *thiophenivorans* DSM 7216^T^, *Ochrobactrum* *pseudogrignonense* CCUG 30717^T^, *Ochrobactrum* *pituitosum* CCUG 50899^T^, *Ochrobactrum* *grignonense* OgA9a^T^, *Ochrobactrum* *rhizosphaerae* PR17^T^, *Ochrobactrum* *anthropi* ATCC 49188^T^, *Ochrobactrum* *lupini* LUP21^T^, *Ochrobactrum* *intermedium* LMG 3301^T^). Information retrieved from MinPath Annotation Tool.

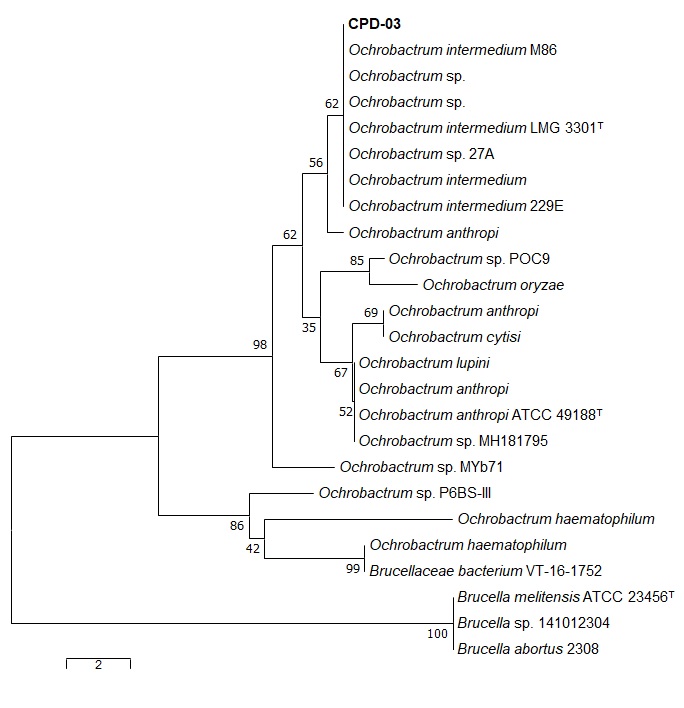

**Supplementary Figure. 3**: Phylogenetic tree with respect to the Arylesterase enzyme present in *Ochrobactrum* strains. These sequences were obtained from UniProt followed by the Top 25 hits were analyzed in MEGA 7.

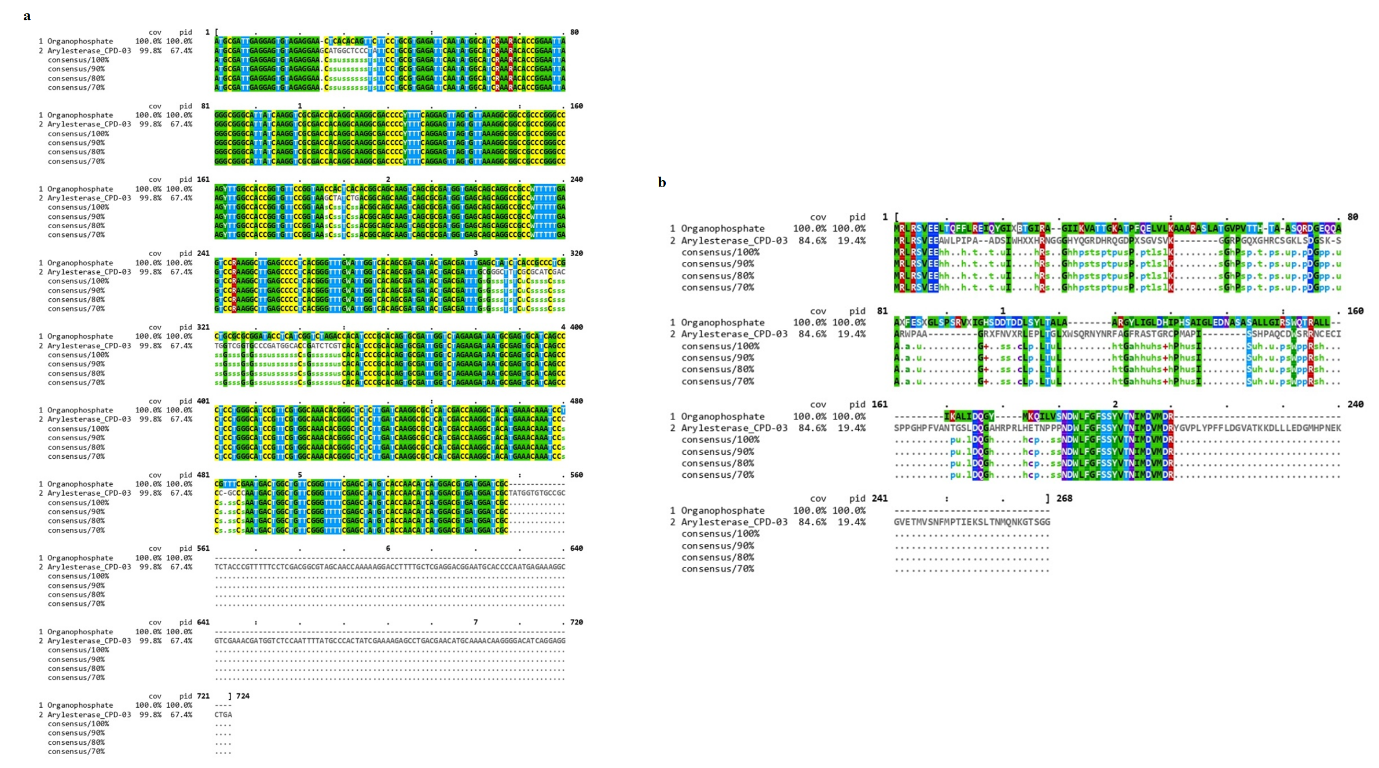

**Supplementary Figure. 4**: a. Nucleotide sequence similarity of organophosphate hydrolase and Arylesterase (EC 3.1.1.2) present in CPD-03 strain; b. Protein sequence similarity of organophosphate hydrolase and Arylesterase (EC 3.1.1.2) present in CPD-03 strain. The nucleotide sequence of organophosphate hydrolase gene was obtained after successful cloning followed by Sangers sequencing and the Arylesterase sequence was obtained from the whole genome. These sequences similarity comparison was performed in ClustalOmega.

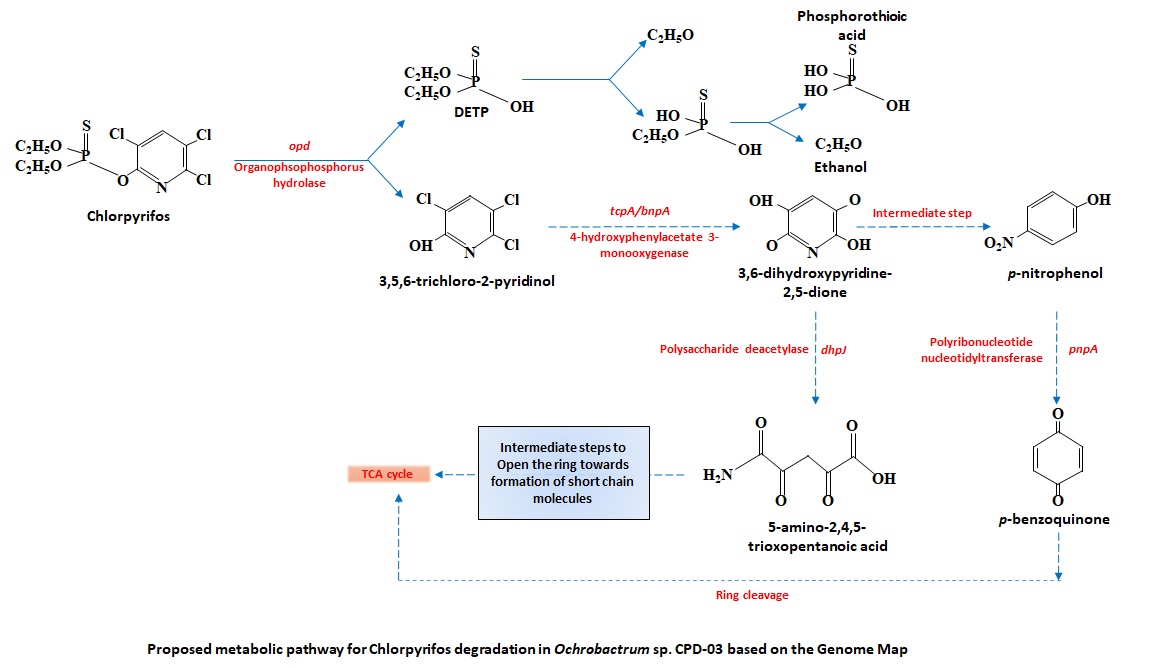

**Supplementary Figure. 5**: Proposed pathway for metabolic degradation of CP by *Ochrobactrum* sp. CPD-03 based on whole genome analysis

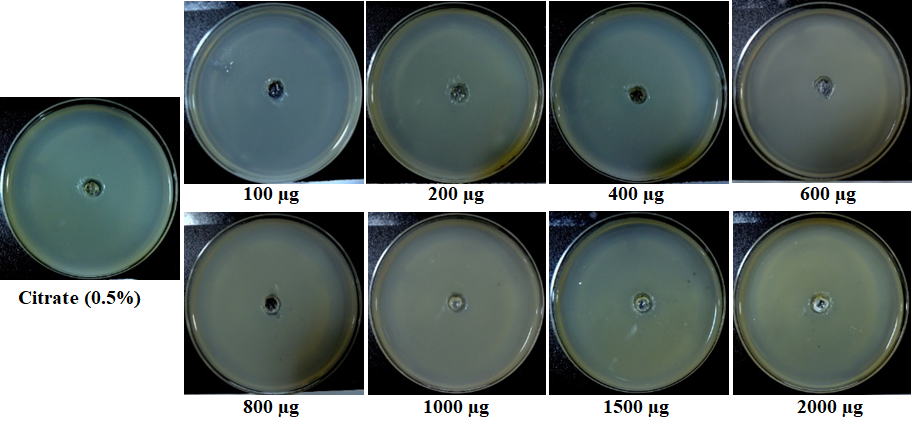

**Supplementary Figure. 6** : Chemotaxis of *Ochrobactrum* sp. CPD-03 resting cells of CPD-03 (1-1.5x10^8^ cells ml^-1^) towards various concentration of CP along with citrate (0.5%) as positive control citrate. Migrated cell width of CPD-03 showing the response towards CP at various concentration

| **Test** | **Results** |
| --- | --- |
| Catalase, Oxidase | + |
| NaCl tolerance | Up to 4% |
| Growth temperature | 25^0^C - 37^0^C (30±2 ^0^C optimum) |
| pH | 5-8 (7 optimum) |
| **Utilization** of  Dextrin, maltose, Cellobiose, Sucrose, N-Acetyl-D-Glucosamine, Citrate, Galactose, Glucose, Arabitol, myo-Inositol, Mannose, Fructose, Xylose | + |

Notes.: (+, positive reaction). Data taken from GEN III plate of Biolog^®^ System

| **Assimilation of** | A | B | C |
| --- | --- | --- | --- |
| Tween 80 | - | + | - |
| N-Acetyl-D-galactosamine | + | + | + |
| Adonitol | + | + | + |
| D-Arabitol | + | + | + |
| Cellobiose | + | + | + |
| Gentiobiose | + | + | + |
| L-Fucose | + | + | + |
| myo-Inositol | + | + | + |
| D-Mannitol | + | + | + |
| L-Rhamnose | + | + | - |
| D-Sorbitol | + | + | + |
| Raffinose | - | - | + |
| Sucrose | + | + | + |
| Trehalose | - | - | + |
| Citric acid | + | + | + |
| Glycerol | - | - | - |

**Supplementary Table**. 1a Phenotypic characteristics of the strain *Ochrobactrum* sp. CPD-03, 1b. Differential biochemical properties of strain CPD-03 (A) and closet reference *Ochrobactrum* type strains: (B) Ca-34^T^; (C) LMG 3301^T^. The table is arranged according to phylogenetic clustering. All data for CPD-03 are from this study and others obtained from Imran et al., 2010.

| RT | Response | Ar/Ht | RFact | ECL | Peak Name | Percent |
| --- | --- | --- | --- | --- | --- | --- |
| 1.4094 | 1143 | 0.011 | 1.180 | 10.6984 | 11:0 anteiso | 1.01 |
| 1.4701 | 1393 | 0.016 | 1.153 | 11.0051 | 11:0 | 1.20 |
| 1.5098 | 459 | 0.009 | 1.137 | 11.1740 | 10:0 2OH | 0.39 |
| 1.6360 | 833 | 0.009 | 1.094 | 11.7107 | 12:0 anteiso | 0.68 |
| 1.8971 | 849 | 0.014 | 1.027 | 12.7126 | 13:0 anteiso | 0.65 |
| 2.1184 | 628 | 0.014 | 0.987 | 13.4833 | 12:0 3OH | 0.46 |
| 2.2721 | 782 | 0.015 | 0.965 | 14.0005 | 14:0 | 0.56 |
| 2.8562 | 1163 | 0.010 | 0.912 | 15.8393 | Sum In Feature 3 | 0.79 |
| 2.9082 | 8304 | 0.008 | 0.910 | 16.0001 | 16:0 | 5.66 |
| 3.0882 | 1047 | 0.013 | 0.902 | 16.5564 | 17:1 anteiso w9c | 0.71 |
| 3.1724 | 725 | 0.012 | 0.899 | 16.8162 | 17:1 w8c | 0.49 |
| 3.1956 | 379 | 0.006 | 0.899 | 16.8879 | 17:1 w6c | 0.25 |
| 3.2028 | 545 | 0.008 | 0.898 | 16.9102 | 17:0 cyclo | 0.37 |
| 3.2327 | 1643 | 0.009 | 0.898 | 17.0024 | 17:0 | 1.10 |
| 3.4100 | 804 | 0.012 | 0.894 | 17.5528 | 16:0 3OH | 0.54 |
| 3.5035 | 59266 | 0.009 | 0.893 | 17.8430 | Sum In Feature 8 | 39.64 |
| 3.5342 | 582 | 0.009 | 0.893 | 17.9382 | 18:1 w5c | 0.39 |
| 3.5529 | 10971 | 0.009 | 0.893 | 17.9963 | 18:0 | 7.34 |
| 3.5797 | 1234 | 0.011 | 0.893 | 18.0812 | 18:1 w7c 11-methyl | 0.83 |
| 3.7517 | 1304 | 0.019 | 0.893 | 18.6277 | 19:0 iso | 0.87 |
| 3.8096 | 1653 | 0.017 | ---- | 18.8116 | unknown 18.815 | ---- |
| 3.8446 | 41251 | 0.009 | 0.894 | 18.9225 | 19:0 cyclo w8c | 27.61 |
| 3.8695 | 998 | 0.015 | 0.894 | 19.0015 | 19:0 | 0.67 |
| 3.9118 | 5990 | 0.011 | 0.895 | 19.1386 | 18:1 2OH | 4.01 |
| 4.0464 | 2199 | 0.010 | 0.897 | 19.5748 | 18:0 3OH | 1.48 |
| 4.0939 | 1558 | 0.013 | 0.898 | 19.7286 | 20:2 w6,9c | 1.05 |
| 4.1292 | 1844 | 0.011 | 0.898 | 19.8430 | 20:1 w7c | 1.24 |
| ---- | 1163 | --- | ---- | ---- | Summed Feature 3 | 0.79 |
| ---- | 59266 | --- | ---- | ---- | Summed Feature 8 | 39.64 |

**Supplementary Table 1c**.; Fatty Acid Methyl Ester (FAME) analysis of CPD-03

| S No. | Compound Name | Gene Designation | KEGG Number | EC Number | Enzyme Name |
| --- | --- | --- | --- | --- | --- |
| 1 | Benzoate degradation | *pobA* | K00481 | 1.14.13.2 | p-hydroxybenzoate 3-monooxygenase |
|  |  | *pcaG* | K00448 | 1.13.11.3 | protocatechuate 3,4-dioxygenase, alpha subunit |
|  |  | *pcaB* | K01857 | 5.5.1.2 | 3-carboxy-cis,cis-muconate cycloisomerase |
|  |  | *pcaC* | K01607 | [4.1.1.44](https://www.kegg.jp/dbget-bin/www_bget?ec:4.1.1.44) | 4-carboxymuconolactone decarboxylase |
|  |  | *dmpB, xylE* | K00446 | 1.13.11.2 | [catechol 2,3-dioxygenase](https://www.kegg.jp/dbget-bin/www_bget?ec:1.13.11.2) |
|  |  | *pcaD* | K01055 | 3.1.1.24 | 3-oxoadipate enol-lactonase |
|  |  | *fadA, fadI* | K00632 | 2.3.1.16 | acetyl-CoA acyltransferase |
|  |  | *dmpB, xylE* | K00446 | 1.13.11.2 | catechol 2,3-dioxygenase |
|  |  | *fadJ* | K01782 | 1.1.1.35/ 4.2.1.17/ 5.1.2.3 | 3-hydroxyacyl-CoA dehydrogenase / enoyl-CoA hydratase / 3-hydroxybutyryl-CoA epimerase |
|  |  | *fadA, fadI* | K00632 | 2.3.1.16 | acetyl-CoA acyltransferase |
|  |  | *gcdH* | K00252 | 1.3.8.6 | glutaryl-CoA dehydrogenase |
|  |  | *paaF, echA* | K01692 | 4.2.1.17 | enoyl-CoA hydratase |
|  |  | *paaH, hbd, fadB, mmgB* | K00074 | 1.1.1.157 | 3-hydroxybutyryl-CoA dehydrogenase |
|  |  | *atoB* | K00626 | 2.3.1.9 | acetyl-CoA C-acetyltransferase |
| 2 | Aminobenzoate degradation | *amiE* | K01426 | 3.5.1.4 | amidase |
|  |  | *bsdC* | K16239 | 4.1.1.61 | 4-hydroxybenzoate decarboxylase subunit C |
|  |  | *paaF, echA* | K01692 | 4.2.1.17 | enoyl-CoA hydratase |
| 3 | Chloroalkane and chloroalkene degradation | *adhP* | K13953 | 1.1.1.1 | alcohol dehydrogenase, propanol-preferring |
|  |  | *aldH* | K00128 | 1.2.1.3 | aldehyde dehydrogenase (NAD+) |
|  |  | *fdhA* | K00148 | 1.2.1.46 | glutathione-independent formaldehyde dehydrogenase |
|  |  | NA | K01560 | 3.8.1.2 | 2-haloacid dehalogenase |
| 4 | Chlorocyclohexane and chlorobenzene degradation | *dmpB, xylE* | K00446 | 1.13.11.2 | catechol 2,3-dioxygenase |
|  |  | NA | K01560 | 3.8.1.2 | 2-haloacid dehalogenase |
| 5 | Xylene degradation | *dmpB, xylE* | K00446 | 1.13.11.2 | catechol 2,3-dioxygenase |
| 6 | Nitrotoluene degradation | *nfsA* | K10678 | 1.5.1.34 | nitroreductase |
| 7 | Ethylbenzene degradation | *fadA, fadI* | K00632 | 2.3.1.16 | acetyl-CoA acyltransferase |
| 8 | Styrene degradation | *amiE* | K01426 | 3.5.1.4 | amidase |
|  |  | *dmpB, xylE* | K00446 | 1.13.11.2 | catechol 2,3-dioxygenase |
| 9 | Atrazine degradation | *ure* | K01427 | 3.5.1.5 | urease |
| 10 | Dioxin degradation | NA | K00480 | 1.14.13.1 | salicylate hydroxylase |
| 11 | Naphthalene degradation | NA | K00480 | 1.14.13.1 | salicylate hydroxylase |
|  |  | *adhP* | K13953 | 1.1.1.1 | alcohol dehydrogenase, propanol-preferring |
| 12 | Polycyclic aromatic hydrocarbon degradation | NA | K00480 | 1.14.13.1 | salicylate hydroxylase |
|  |  | *pcaG* | K00448 | 1.13.11.3 | protocatechuate 3,4-dioxygenase, alpha subunit |

**NA*: Not assigned in the genome of CPD-03**

**Supplementary Table 2**. Putative genes of *Ochrobactrum* CPD-03 involved in degradation of Aromatic and relative compounds. Information retrieved from KAAS annotation tool. [NA: Not available]

| **S No.** | **Activity** | **Gene Designation** | **Enzyme Name** |
| --- | --- | --- | --- |
| 1 | IAA production | **NA*** | Indole-3-glycerol phosphate synthase |
| 2 | Siderophore production | *FhuB* | Ferric hydroxamate ABC transporter, permease component |
|  |  | *TonB* | Ferric siderophore transport system, periplasmic binding protein |
|  |  | *FepC* | Ferric enterobactin transport ATP-binding protein @ ABC-type Fe3^+^-siderophore transport system, ATPase component |
|  |  | *FepG* | Ferric enterobactin transport system permease protein |
|  |  | *FepB* | Ferric enterobactin-binding periplasmic protein |
|  |  | *FepC* | Ferric enterobactin transport ATP-binding protein |
|  |  | *FhuD* | Ferric hydroxamate ABC transporter, periplasmic substrate binding protein |
|  |  | *FhuC* | Ferric hydroxamate ABC transporter, ATP-binding protein |
|  |  | **NA*** | ABC transporter (iron.B12.siderophore.hemin) , permease component |
|  |  | *FhuA* | Ferric hydroxamate outer membrane receptor |
|  |  | *EntS* | Enterobactin exporter |
|  |  | **NA*** | ABC-type Fe3+-siderophore transport system, permease component |
|  |  | *FhuB* | Ferric hydroxamate ABC transporter, permease component |
| 3 | Flagellar protein | *FlhB*, *FliL*, *FliP*, *FliQ*, *FlhA*, *FliR* | Flagellar biosynthesis protein |
|  |  | *FliG*, *FliN*, *FliM* | Flagellar motor switch protein |
|  |  | *MotA*, *MotB* | Flagellar motor rotation protein |
|  |  | *FlgF*, *FlgF* | Flagellar basal-body rod protein |
|  |  | *FliI* | Flagellum-specific ATP synthase |
|  |  | *FlgB*, *FlgC*, *FlgG* | Flagellar basal-body rod protein |
|  |  | *FliE* | Flagellar hook-basal body complex protein |
|  |  | *FlgA* | Flagellar basal-body P-ring formation protein |
|  |  | *FlgI* | Flagellar P-ring protein |
|  |  | *FlgH* | Flagellar L-ring protein |
|  |  | *FlaA*, *FlaF*, *FliJ* | Flagellin protein |
|  |  | *FliF* | Flagellar M-ring protein |
|  |  | *FliK* | Flagellar hook-length control protein |
|  |  | *FlgE* | Flagellar hook protein |
|  |  | *FlgK*, *FlgL* | Flagellar hook-associated protein |
|  |  | *FlgD* | Flagellar basal-body rod modification protein |
|  |  | *FlgJ* | Flagellar protein [peptidoglycan hydrolase] |
| 4 | Chemotaxis | **NA*** | Dipeptide-binding ABC transporter, periplasmic substrate-binding component |
|  |  | *RbsB* | Ribose ABC transporter, periplasmic ribose-binding protein |
|  |  | *MalE* | Maltose/maltodextrin ABC transporter, substrate binding periplasmic protein |
|  |  | **NA*** | Dipeptide-binding ABC transporter, periplasmic substrate-binding component; Putative hemin-binding lipoprotein |
|  |  | *CheR* | Chemotaxis protein methyltransferase (EC 2.1.1.80) |
|  |  | *motC* | Chemotaxis protein |
|  |  | *MalE* | Maltose/maltodextrin ABC transporter, substrate binding periplasmic protein |
|  |  |  | Methyl-accepting chemotaxis protein |
| 5 | HCN production, Phosphate solubilization, ACC deaminase activity, Ammonia production | **NA*** | |

**Supplementary Table** **3**. Putative genes of *Ochrobactrum* CPD-03 involved in Plant growth promoting activities and chemotaxis. Information retrieved from RAST annotation tool. [NA: Not available]
